# Toxin resistance in the monarch butterfly expanded its milkweed host range

**DOI:** 10.64898/2026.05.25.727632

**Authors:** Naama C. Weksler, Michael Astourian, Derrick Yip, Elizabeth Ordeman, Kirsten Verster, Boris Korablev, Susan L. Bernstein, Anurag A. Agrawal, Noah K. Whiteman, Marianthi Karageorgi

## Abstract

A central problem in ecology and evolution is to understand the extent to which herbivorous insect host plant specialization has been shaped by bottom-up natural selection from host-plant chemical defenses or top-down natural selection from natural enemies. Here, we show that the stepwise evolution of ATPα target-site insensitivity (TSI) in the monarch butterfly lineage consistently tracked a dietary association with cardenolide-defended *Asclepias* host plants. We further show that *Drosophila melanogaster* flies edited with CRISPR-Cas9 genome engineering to carry the monarch butterfly’s more derived *ATPα* genotype maintain survival across *Asclepias* host-plant diets that span a broad cardenolide defense gradient, whereas flies carrying early-diverging *ATPα* genotypes in the monarch lineage show lower survival. We conclude that a resistance trait linked to sequestration and enemy protection also expanded dietary resistance to highly defended host plants, revealing how a single molecular adaptation may have been driven by both top-down and bottom-up selective pressures during herbivore specialization.

## Main text

Toxin resistance in specialist herbivores can be shaped by both bottom-up pressures from plant chemical defenses and top-down pressures from natural enemies when plant toxins are sequestered for protection. These defenses impose barriers that specialist herbivores must overcome to exploit defended host plants ^1^. However, these toxins can also be sequestered and deployed against predators and parasitoids ^2^, raising the possibility that adaptive traits shaped by one set of selective agents can also enable adaptation to others. Disentangling the adaptive roles of toxin resistance in herbivorous insects is challenging because the same trait can both confer dietary resistance to plant defenses and prevent toxicity following toxin uptake and sequestration.

The tribe Danaini (Danainae, Nymphalidae), commonly known as milkweed butterflies, provides a powerful system for studying how toxin resistance mediates both host use and enemy protection. Milkweed butterflies frequently use cardenolide-defended host plants ^3^, especially milkweeds (Apocynaceae), and monarch butterflies (*Danaus plexippus*) feed on cardenolide-defended *Asclepias* hosts ^4^ while sequestering these toxins for enemy defense ^5^. Cardenolides inhibit the Na⁺,K⁺-ATPase, and resistance in the monarch lineage evolved through substitutions in the first extracellular loop of ATPα at positions 111, 119 and 122, producing stepwise increases in target-site insensitivity (TSI) ^6,7^. Previous research found that the monarch ATPα state confers extreme TSI and is associated with sequestration and protection from toxicity ^6,8^. Whether this extreme ATPα state also increases survival on host-plant diets spanning a natural cardenolide defense gradient, beyond earlier ATPα states, has remained unclear.

Here, we first addressed how the evolution of *Asclepias* host use related to stepwise increased ATPα TSI across Danaini. To infer the evolutionary history of *Asclepias* use, we reconstructed host-use evolution across Danaini by generating a species-level molecular phylogeny and curating larval host-plant records ^9^ together with butterfly and host-plant taxonomy (see **Methods**). We focused on 55 of approximately 160 Danaini species for which both phylogenetic placement and host-use data were available. Host use within the milkweed clade of Apocynaceae, which comprises the subfamilies Asclepiadoideae and Secamonoideae ^10^, showed significant phylogenetic signal, indicating that closely related Danaini species tended to share milkweed-clade associations (Pagel’s λ = 0.58, LR = 22.32, *P* = 2.3 × 10-6; **Supplementary Fig. 1**).

Ancestral state reconstruction further revealed an evolutionary shift from broader milkweed-clade use to *Asclepias* use within *Danaus* (maximum likelihood model with symmetric transition rates, see **Methods**) (**Fig. 1**). Association with the broader milkweed clade was inferred in the tribes Danaina and Amaurina (marginal state probability = 0.77 and 0.83, respectively), followed by strong support for *Asclepias* association at the base of the *Danaus* lineage (marginal state probability = 0.99). Within *Danaus*, support for association with *Asclepias* was strongest in the subgenera *Danaus* and *Anosia* (marginal state probability = 0.98 and 0.97, respectively) ^11^, both of which include lineages distributed within the native range of *Asclepias* in the Americas. Support was lower in *Salatura*, a primarily Oriental–Australasian subgenus ^11^ largely outside the native range of *Asclepias* (marginal state probability = 0.78).

**Figure 1.**
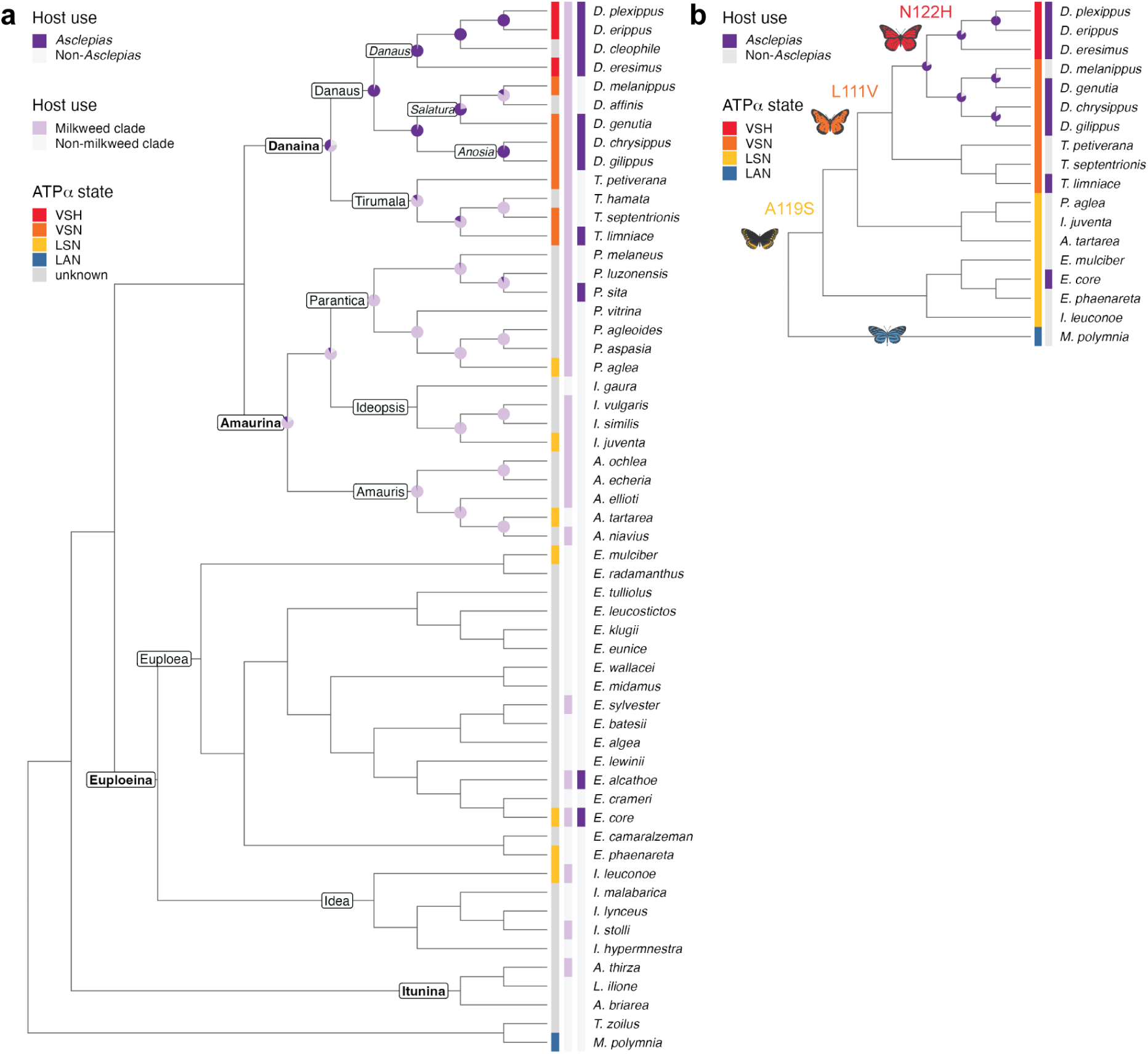
Consistent *Asclepias* use in *Danaus* coincided with the stepwise evolution of extreme target-site insensitivity to cardenolides in ATPα. **a,** Maximum-likelihood phylogeny of 55 Danaini butterflies and two outgroups within the Danainae subfamily with available host-plant data, pruned from a tree inferred in this study from a 6,585-bp concatenated alignment of mitochondrial *COI* and 11 nuclear gene regions (see **Methods**). Danaini subtribes, genera, and *Danaus* subgenera are annotated on the tree. Of note, *D. eresimus* is treated with the *Danaus* subgenus rather than *Anosia* because it clusters with the *Danaus* subgenus in our tree, consistent with Aardema & Andolfatto, 2016 ^12^. Pie charts show maximum-likelihood-based marginal probabilities of ancestral host-plant associations with non-milkweed hosts (grey), the milkweed clade excluding *Asclepias* (light purple), and the *Asclepias* genus (dark purple). Pie charts are shown only for nodes at which the marginal probability of either milkweed-clade association (calculated as the sum of the milkweed clade and *Asclepias* states) or *Asclepias* association is greater than 0.75. The adjacent heatmaps indicate documented larval host associations with the milkweed clade and the genus *Asclepias*, alongside ATPα states associated with increasing Na⁺,K⁺-ATPase insensitivity to cardenolides (LAN, LSN, VSN, VSH; unknown in grey). Presence was defined as at least one documented larval host association within the corresponding plant lineage, whereas absence indicates no recorded association. See also the complete plant–butterfly host-use matrix in **Supplementary Fig. 1**. **b**, Subset of phylogeny from (**a**) filtered for species for which ATPα state is known. Inferred transitions at ATPα sites 111, 119, and 122 are mapped at relevant nodes according to ^6^ and validated using maximum likelihood and parsimony-based ASR.

The inferred origins of three different ATPα states conferring stepwise increases in TSI correlated with this host-use transition within Danaini. ATPα state transitions were mapped to the 55-species Danaini tree based on previous research ^6^ and validated using both maximum likelihood and parsimony-based ASR. The ATPα LSN state, the first step in this trajectory that conferred lower TSI, was inferred to arise at the base of the Euploeina and Amaurina tribes, which were not inferred to associate with *Asclepias*. The ATPα VSN state, the second step that conferred intermediate TSI, was inferred to evolve at the base of the Danaina tribe and includes both lineages inferred to associate with *Asclepias* and lineages not inferred to do so. The ATPα VSH state, which confers the highest TSI, was inferred to evolve in the subgenus *Danaus Danaus*, where *Asclepias* association was consistently inferred. Thus, we conclude from these results that the stepwise evolution of ATPα states culminating in extreme TSI tracks consistent *Asclepias* use, a pattern that was further supported by a geographically filtered reconstruction excluding host records that may reflect introduced *Asclepias* outside the plant’s native range in the Americas (see **Methods**, **Supplementary Fig. 2**).

This phylogenetic association led us to test whether the more derived monarch ATPα VSH state supports survival across *Asclepias* host-plant diets spanning a broad cardenolide defense gradient, relative to more ancestral ATPα states. We compared larval-to-adult survival among edited *D. melanogaster* lines carrying precise amino acid substitutions at sites 111, 119 and 122. These lines represent stepwise accumulation of ATPα substitutions in the monarch lineage (LSN, VSN and VSH), an outgroup Danainae state (LAN), and a cardenolide-sensitive control state (QAN) ^6^. We backcrossed all lines into a common *w1118* background (see **Methods**) and assayed them under the same conditions, allowing us to isolate the contribution of ATPα from other butterfly-specific resistance mechanisms. Larvae were reared on media supplemented with air-dried leaf powder from six *Asclepias* species that are documented monarch hosts ^4,13^ and span a broad gradient of foliar cardenolide defenses ^14^. Unsupplemented media and media supplemented with lettuce (*Lactuca sativa*), which is not known to contain cardenolides, served as non-cardenolide controls (see **Methods**, **Fig. 2a**).

**Figure 2.**
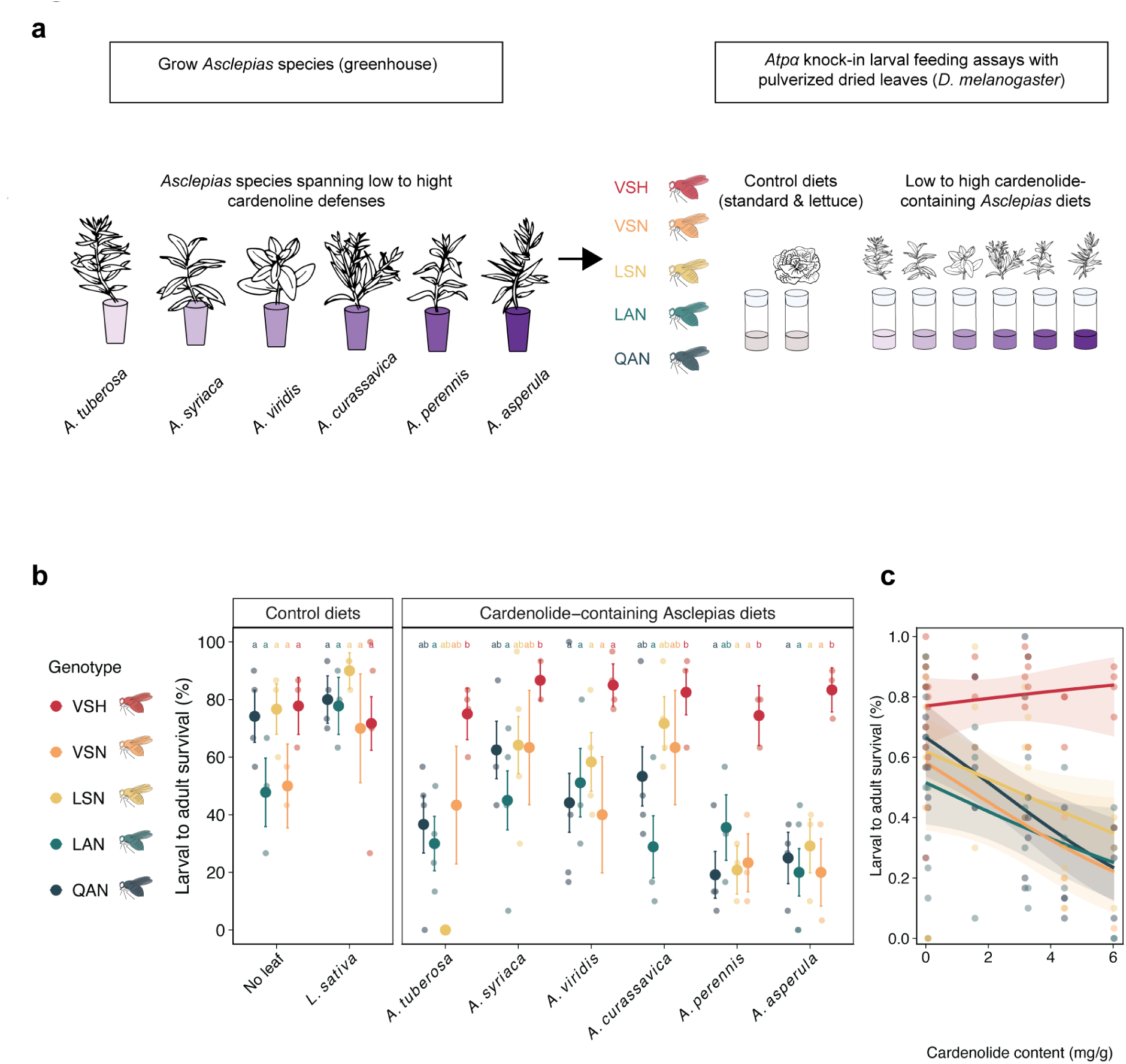
The monarch *ATPα* VSH genotype conferred higher survival across increasingly cardenolide-defended *Asclepias* diets relative to other Danaini and outgroup *ATPα* genotypes. **a**, Six *Asclepias* species spanning a gradient of cardenolide defenses ^14^ were grown in a greenhouse. Harvested leaves were dried, then ground into powder, and incorporated into standard Drosophila media. Larval-to-adult survival of knock-in *Drosophila melanogaster* lines carrying ancestral (QAN), intermediate (LAN, LSN, VSN), and monarch (VSH) *Atpα* genotypes was compared on cardenolide-containing media (supplemented with *Asclepias* leaf material) versus non-cardenolide (standard and supplemented with lettuce leaf material) controls. **b**, Larval-adult fly survival on cardenolide-containing vs control diets (n = 1–4; 30 larvae per replicate; quasibinomial logistic regression; model-estimated mean ± s.e). Letters indicate groups from Bonferroni-adjusted pairwise post hoc comparisons. **c**, Larval-to-adult survival of flies as a function of approximate cardenolide concentration present in leaves of the different milkweed species ^14^ (quasi-binomial logistic regression; mean ± c.i.).

Fly survival differed among *Atpα* genotypes on *Asclepias* diets but not on non-cardenolide controls (φ = 6.7, quasi-binomial logistic regression) (**Fig. 2b**, **Supplementary Table S1**). Across *Asclepias* diets, flies carrying the monarch VSH genotype had 2.0–2.4-fold higher model-estimated survival than QAN, LAN, LSN or VSN flies (all Bonferroni-adjusted pairwise *P* ≤ 5.9 × 10^-6^). By contrast, survival did not differ between fly genotypes on media not containing cardenolides (all pairwise *P* ≥ 0.59), indicating that the VSH advantage was specific to defended milkweed diets rather than reflecting differences in baseline viability.

Diet-specific comparisons showed that the VSH survival advantage was most pronounced on the most highly defended diets (**Supplementary Table S2**). On *A. asperula*, the species with the highest foliar cardenolide concentration in our panel, VSH flies survived better than all other genotypes. On *A. perennis*, the second-highest cardenolide diet, VSH survival exceeded QAN, LSN and VSN, but not LAN. Among lower- to intermediate-cardenolide diets, VSH also exceeded LAN on *A. tuberosa*, *A. syriaca* and *A. curassavica* (Bonferroni-adjusted pairwise *P* = 0.029, 0.047, and 0.012, respectively). No consistent survival differences were detected among the non-VSH genotypes on any diet.

We next addressed whether genotype-specific survival changed across the *Asclepias* cardenolide defense gradient. Using published foliar cardenolide concentration estimates as a proxy for defense level ^14^, we found that only VSH did not decline in survival as cardenolide concentration increased (b = 0.07 ± 0.10, z = 0.69, *P* = 0.48) (**Fig. 2c**, **Supplementary Table S3**). In contrast, QAN, LAN, LSN and VSN all had negative slope estimates across the gradient, with significant declines for QAN, LAN and LSN (b = −0.31 to −0.18; *P* ≤ 0.044) and a similar negative trend for VSN (b = −0.26, *P* = 0.057). Pairwise slopes comparisons showed the strongest contrast between VSH and the sensitive QAN control (*P* = 0.028), consistent with previous evidence for stepwise increases in ATPα TSI across the monarch-lineage genotypes ^6^.

Foliar cardenolide concentration also captured the broader *Asclepias* defense gradient in this diet panel because it covaried with cardenolide diversity (r = 0.85) and polarity (r = −0.71), two additional axes of cardenolide defenses ^14^. Under dried whole-leaf assay conditions, VSH survival was maintained along this gradient, indicating that the monarch *ATPα* genotype supported survival across increasingly defended *Asclepias* diets. Although drying and grinding removed physical defenses such as latex exudation, they may retain non-cardenolide secondary metabolites, including phenolics ^15^. Such compounds, whether reduced during processing or tolerated after ingestion through endogenous detoxification mechanisms ^16^ in *D. melanogaster*, did not prevent VSH flies from maintaining survival across the diet panel.

Together, we found that consistent *Asclepias* use in *Danaus* lineages coincided with the ATPα VSH state, and that this state was sufficient in a novel insect genomic background to maintain high relative survival across a host-plant defense gradient, including highly defended milkweeds. The ATPα VSH state is restricted to the *Danaus* subgenus, whose range partially overlaps the native radiation of *Asclepias* in North America ^11,17^, consistent with a geographic concordance between extreme ATPα TSI and *Asclepias* host use. The sporadic occurrence of this host association across milkweed butterflies (see **Supplementary Fig. 1**) further suggests that other resistance mechanisms can also support *Asclepias* use. Some milkweed butterflies carrying less resistant LSN or VSN ATPα states have documented *Asclepias* host records ^9^ and cope with highly defended *Asclepias* species ^8^, although some of these records may reflect recent associations with introduced *Asclepias* outside the plant’s native range (see **Supplementary Fig. 2**). Thus, the evolution of extreme ATPα TSI in the *Danaus Danaus* subgenus may provide one route to dietary resistance, while additional mechanisms, such as efflux transporters ^18,19^, also enable feeding on chemically defended *Asclepias* host plants.

Overall, our study shows that a molecular resistance trait previously linked to sequestration and enemy protection ^6,8^ can also enhance dietary resistance to defended host plants. This illustrates how a single adaptation may have been shaped by both top-down and bottom-up selective pressures during herbivore specialization. In the monarch butterfly lineage, the stepwise evolution of extreme ATPα TSI may have initially reduced the costs of toxicity that are associated with cardenolide sequestration, consistent with the independent evolution of VSH in the sequestering leaf beetle *Chrysochus cobaltinus* ^20^. By increasing tolerance to highly defended milkweed diets, this same molecular adaptation may also have enabled monarchs to exploit *Asclepias* host plants across regions where monarch butterflies and milkweeds co-occur. This is consistent with the hypothesis that key innovations can unlock access to previously unavailable resources ^21^.

## Methods

### Phylogenetic signal and ancestral reconstruction of host-plant use in Danaini

To test whether host-plant use is phylogenetically conserved and to reconstruct the evolutionary history of milkweed-clade association in Danaini butterflies, we combined a species-level phylogeny of Danaini with curated larval host-plant records and genus-level phylogenies of Apocynaceae and Moraceae. We analyzed a subset of 55 Danaini species for which both phylogenetic placement and larval host-plant data were available. Phylogenetic signal in milkweed-clade association was quantified using Pagel’s λ, and ancestral state reconstruction was used to infer the evolutionary origin of milkweed-clade and *Asclepias* use.

#### Danaini species phylogeny

To infer a maximum-likelihood phylogeny of the tribe Danaini, we used DNA sequence data from 73 Danaini species and two outgroup species, *Tellervo zoilus* (Tellervini) and *Mechanitis polymnia* (Ithomiini), for which at least one of the following 12 gene regions was available in GenBank (February 2026): the mitochondrial *COI* gene and the nuclear genes *ArgKin*, *CAD*, *DDC*, *EF1α*, *GAPDH*, *IDH*, *MDH*, *RpS5*, *RpS2*, *wingless*, and *ATPα*. These loci overlap with gene sets previously employed in Danaini phylogenetic studies ^22,23^, ensuring comparability with previous analyses of the tribe (see below for details).

We assembled our dataset using the R package rentrez (v.1.2.4) to retrieve all available sequences for each locus and species from GenBank. For each gene, accessions were filtered using locus-specific length thresholds to exclude short fragments. When multiple sequences were available for a given species and locus, up to three longest accessions were retained for alignment. Nuclear protein-coding genes were translated and aligned at the amino acid level using MAFFT and back-translated to nucleotides to preserve codon structure, whereas COI was aligned using the invertebrate mitochondrial genetic code. Alignments were manually inspected in Galaxy (https://usegalaxy.eu) to identify short or misannotated sequences; problematic accessions were removed and alignments were re-run iteratively. Final alignments were trimmed to homologous codon-consistent regions to preserve reading frame and ensure correct codon-position partitioning. For each species and locus, the sequence with the longest overlap within the trimmed alignment was retained. Species names were standardized across loci prior to concatenation. Trimmed loci were concatenated into a supermatrix of 6,585 base pairs across 75 taxa, with missing loci represented as gap-only blocks. Partition coordinates were defined for each locus and subdivided by codon position (1st, 2nd, and 3rd positions per locus). Maximum-likelihood inference was performed in IQ-TREE v2 using ModelFinder with partition merging (–m MFP+MERGE) and 1,000 ultrafast bootstrap replicates (–B 1000, –bnni). ModelFinder selected a partitioned model under BIC, merging the codon-position partitions into six subsets with separate substitution models. The best maximum-likelihood tree had a log-likelihood of −32,509.8833. The resulting tree was rooted using the two outgroup taxa. The GenBank accession numbers of all sequences used for phylogenetic inference are provided in **Suppl. Data 1**.

#### Host-plant association data

We analyzed published larval hostplant records compiled by Ackery ^9^, encompassing 68 of approximately 160 described Danaini species. We also added data for three Danaini species not included in Ackery ^9^ from the HOSTS database ^24^. These host-use data were integrated with the current taxonomic classification of Danaini butterflies ^22^ and the current taxonomic classification of their primary cardenolide-containing hostplant families Apocynaceae and Moraceae.

Historical plant genus names from hostplant records ^9^ were assigned to their current taxonomic positions following ^25,26^, and genus names were recorded based on current synonyms using Plants of the World Online (POWO). In addition, we compiled published information on the occurrence of cardenolides across genera of Apocynaceae and Moraceae from the literature ^27,28^. The full host plant association dataset used for all analyses is provided in **Suppl. Data 2**. We used the Apocynaceae genus-level phylogeny published by Fishbein et al. ^29^. Relationships among Moraceae genera were obtained from a genus-level time-calibrated phylogeny from TimeTree ^30^. Both trees were pruned to include only genera represented in our host-plant dataset.

The Danaini phylogeny included 73 species, and larval host-plant data were available for 71 species. The overlap between the phylogenetic and host-plant datasets comprised 55 species. Phylogenetic signal analyses and ancestral state reconstructions were therefore conducted on this overlapping subset of taxa.

#### Phylogenetic signal and ancestral state reconstruction

We quantified phylogenetic signal in association with the milkweed clade of Apocynaceae (subfamilies Asclepiadoideae and Secamonoideae) using Pagel’s λ ^31^, estimated by maximum likelihood and implemented in the R package phytools (v 2.5.2) ^32^. Milkweed association was coded as a binary trait for each species in the Danaini phylogeny, where species were assigned a value of 1 if at least one documented larval host record belonged to the milkweed clade (subfamilies Secamonoideae or Asclepiadoideae), and 0 otherwise. Host associations were treated as presence-only records, such that any documented larval use of a milkweed-clade host was considered evidence of association. Statistical support for phylogenetic signal was assessed by likelihood-ratio comparison of the fitted model against a model with λ constrained to zero, corresponding to phylogenetic independence of trait values.

We performed ancestral state reconstruction (ASR) of host-plant associations on the Danaini using a discrete-character maximum likelihood framework implemented in the fitMk function of the R package phytools (v. 2.5.2). We modeled host use as a multi-state character with three levels: *Asclepias*, milkweed clade (subfamilies Secamonoideae and Asclepiadoideae), and none (non-milkweed hosts). Host use was coded as a binary trait indicating the presence or absence of larval feeding on each focal plant lineage, and species were assigned to the most specific host category for which feeding was recorded, prioritizing *Asclepias* use followed by general milkweed clade use. Character evolution was modeled using a symmetric (SYM) continuous-time Markov model, allowing for transitions between all states while maintaining transition-rate parsimony. We estimated and visualized marginal ancestral state probabilities for all internal nodes as full likelihood proportions, providing a complete representation of evolutionary uncertainty across the phylogeny.

To account for possible effects of introduced host plants, we performed an additional geographically filtered reconstruction of *Asclepias* association in the focused ATPα dataset. Native geographic ranges of butterfly species were coded from Ackery and Vane-Wright ^3^, and species were scored as having native-range *Asclepias* association only when documented *Asclepias* host use occurred in species whose native range overlaps the native range of *Asclepias* in the Americas. Documented *Asclepias* records outside this native-range overlap were retained in the host-use dataset but scored as non-native-range associations in this sensitivity analysis. Ancestral state reconstruction was then repeated using the same maximum-likelihood Mk as described above.

Ancestral ATPα H1–H2 loop sequences were inferred using maximum-likelihood ancestral sequence reconstruction in the R package ace as well as parsimony-based ancestral state reconstruction in the R package castor. An existing in-frame codon alignment of the 12-amino acid H1-H2 loop was imported from ^6^ and pruned for species in the 55-species Danaini tree. Two species for which sequence data is newly available, *D. eresimus* and *D. melanippus*, were added. Ancestral amino acid sequences were reconstructed at internal nodes and translated to amino acid sequences for visualization.

### Experimental reconstruction of *ATPα* genotypes and survival across the milkweed cardenolide defense gradient

To test the functional consequences of stepwise ATPα evolution, we compared larval to adult survival among *D. melanogaster* knock-in lines carrying reconstructed *ATPα* genotypes corresponding to sequential states in the Danaini lineage and outgroups. All *Atpα* knock-in lines were backcrossed into a common *w1118* background to minimize background effects. To test for resistance to milkweed toxins, larvae were then reared on artificial diets supplemented with dried leaf powder from milkweed species spanning a natural gradient of cardenolide defenses. Survival was quantified from second instar to adult eclosion and analyzed using quasi-binomial logistic regression.

#### Backcrossing *ATPα* knock-in fly lines in *w1118* background

We previously used two rounds of CRISPR–Cas9 genome editing coupled with homology-directed repair (HDR) to generate viable, homozygous *Atpα* knock-in *D. melanogaster* lines carrying the precise substitutions at sites 111, 119 and 122 of 4 consecutive *Atpα* genotypes found in milkweed butterflies (Q111L/A119S, Q111V/A119S, and Q111V/A119S/N122H) as well as the outgroup Q111L found in an outgroup, and in *D. melanogaster*, QAN ^6^. Here, we backcrossed each of these *Atpα* knock-in and control lines for 7 generations into the *w1118* background to eliminate potential off-target mutations from the

CRISPR-Cas9 genome editing ^33^. In the first backcross, homozygous males for the *ATPα* knock-in allele were crossed with *w1118* virgin females. In the second backcross, heterozygous male progeny for the *ATPα* knock-in allele were crossed with *w1118* virgin females. This initial use of males ensured replacement of the X chromosome with the *w1118* background. In subsequent backcrosses, heterozygous virgin female progeny for the *ATPα* knock-in allele were crossed with *w1118* males, taking advantage of recombination in females to further replace the original background with *w1118*. At each backcross, heterozygous female progeny carrying the *ATPα* knock-in alleles were selected by PCR genotyping for the next generation. Finally, heterozygous male and female progeny carrying the *ATPα* knock-in alleles from generation 7 were crossed and homozygous progeny were selected to re-establish homozygous *Atpα* knock-in lines.

To confirm each genotype, we extracted genomic DNA from single midlegs using a proteinase K-based lysis buffer (10mM Tris-Cl pH 8.2, 1mM EDTA, 25mM NaCl, 500μg/ml Proteinase K) with overnight incubation at 56°C followed by enzyme inactivation at 95°C for 5 minutes. We performed PCR amplification using line-specific primers: ATPaF2 (5’-GGGTCTCGCTGCAGTGCATC-3’) and ATPaR2 (5’-CACCTCAACGACATCGCCCAG-3’) for Atpα knock-in lines and 11B_F1 (5’-CATCCCCGTGCACAGGTTCGG-3’) and 11B_R1 (5’-GGCGTGACTTAGACCCTGCGG-3’) for the control engineered line. PCR conditions were 30 cycles of 95°C for 30s, 60°C (55°C for control line) for 30s, and 68°C for 1 minute, with products verified by Sanger sequencing.

#### Milkweed growing and harvesting

We grew and harvested six milkweed species spanning a range of cardenolide defenses : *A. tuberosa*, *A. syriaca*, *A. viridis*, *A. curassavica*, *A. perennis*, and *A. asperula* ^34^. Seeds of *A. tuberosa*, *A. syriaca*, *A. viridis*, *A. curassavica*, and *A. asperula* were purchased from the Joyful Butterfly nursery, while *A. perennis* seeds were obtained from Bradford Grimm/Ginie Anthony. We germinated seeds of all milkweed species using either a cold-stratification protocol (*A. perennis*) developed in the lab of Prof. Anurag Agrawal or a water-germination protocol (all other species) developed by Bradford Grimm. In the water germination protocol, the milkweed seeds were soaked in Milli-Q water in a plastic cup on a 26°C warming mat for 7-14 days. The water was replaced every ∼12 hours to avoid fungal growth. In the cold-stratification protocol, the seeds were treated twice with 5% bleach in Milli-Q water for 3 minutes, scarified and cold stratified in dark at 4°C on moist filter paper for 7 days. Following the cold-stratification, the seeds were soaked in Milli-Q water in the dark at 28°C for 7 days. After germination, the seedlings were planted into flat trays of potting soil (Sun Gro Sunshine All-Purpose Potting Mix). Upon reaching the four-leaf stage, one plant per pot (11cm x 11cm x 24cm) was transplanted into a mix of soil (½ Sunshine mix, ¼ turface, ¼ perlite) with ⅓ teaspoon of slow-release fertilizer (Osmocote Plus Outdoor & Indoor Smart-Release Plant Food) and grown in a greenhouse (12h daylight, over 400uE of light, 26°C day : 20°C night) at the Oxford Research Facility at UC Berkeley. All plants were watered ad libitum once every week.

*Amblyseius cucumeris* was sprinkled on plants every two weeks to avoid thrip infestation. We harvested leaves from non-flowering plants after four to six months of growth. Τhe leaves were cut from the plant at the petiole to stimulate growth of new leaves. Organic *Lactuca sativa* (lettuce) leaves were bought from the North Berkeley Farmer’s market. After harvesting, the leaves were placed into paper envelopes and stored at -20°C in order to lyse cells, thereby liberating cardenolides from the cells. Finally, the leaves were air dried in a drying oven at 40°C/ 20% humidity for three-to-four days. Once dry, the leaves were ground into a fine powder using a Mueller Austria HyperGrind coffee grinder and stored at 4°C until used for experiments. The experimental timeline from milkweed growing to leaf harvesting and pulverization spanned from December 2019 to August 2020.

#### Fly survival on medium supplemented with milkweed leaf powder

We performed all feeding experiments in a walk-in growth chamber at ∼21–23°C and ∼50–60% humidity, on a 12 h:12 h light:dark cycle. In all feeding experiments, vials for independent trials were coded and placed in a randomized order in rows on cardboard trays. We raised flies from the four knock-in (LAN, LSN, VSN, and VSH) and the engineered control (QAN) line on minimal sugar-yeast (SY) Drosophila medium with or without additional pulverized plant material. SY media 35 consisted of 50g sucrose, 100g brewer’s yeast, 15g agar, 3g Nipagin® M and 3ml propionic acid per litre, and was chosen over other common media because it allowed incorporation of plant leaf powder at high concentrations and the fact that use of the SY media alone did not appear to negatively affect fly survival. We introduced 30 second instar larvae of each line in narrow vials containing 5 ml of SY medium^35^, with or without the addition of 20% w/v dried plant leaf powder. We used second-instar larvae for the experiment because of the occurrence of variably penetrant embryonic lethality in the knock-in lines. We selected the minimal SY diet because it enabled successful addition of plant leaf powder to 20% w/v, which more closely mimics the dry mass:water ratio as is found in milkweed foliar tissue, while retaining larval survival in control experiments. Each of the treatment and control groups was tested in one to four replicates. Pupariation, adult eclosion, and survival were monitored over a period of ∼20 days until all adult flies had eclosed.

#### Statistical analysis of fly survival data

To test whether the *ATPα* genotypes differ in their survival response to cardenolide-containing versus non-cardenolide-containing media, we performed a quasi-binomial logistic regression on the probability of survival against model terms for genotype and media type, as well as an interaction term for the effects of genotype and media type combined. We used a quasi-binomial logistic regression model due to data overdispersion. The quasi-binomial logistic regression is of the form log(P(survival)/(1 - P(survival))) = β₀ + β₁(genotype) + β₂(media type) + β₃(genotype × media type), where genotype represents the *ATPα* genotype of each knock-in *D. melanogaster* line and media represents the type (cardenolide-containing or not) of pulverized plant added to the media. Post-hoc pairwise comparisons between genotypes within each media type were conducted using estimated means from the model with Bonferroni correction for multiple testing.

In order to identify which media conditions drove this difference, we performed a quasi-binomial logistic regression on the probability of survival against model terms for genotype and media species, as well as an interaction term for the effects of genotype and media species combined.

As previously, we used a quasi-binomial logistic regression model due to data overdispersion. The quasi-binomial logistic regression is of the form log(P(survival)/(1 - P(survival))) = β₀ + β₁(genotype) + β₂(media species) + β₃(genotype × media species), where media species represents the species of milkweed added to the media. Post-hoc pairwise comparisons between genotypes within each media were conducted using estimated means from the model with Bonferroni correction for multiple testing.

Finally, to test whether the *ATPα* genotypes differed in their survival response to increasing cardenolide concentrations, we performed a quasi-binomial logistic regression on the probability of survival against model terms for genotype and cardenolide concentration, as well as an interaction term for the effects of genotype and cardenolide concentration combined. We used cardenolide concentration measurements for each milkweed species reported in ^14^, and included lettuce (*L. sativa*) as a control with zero cardenolide concentration. We used a quasi-binomial logistic regression model due to data overdispersion. The quasi-binomial logistic regression is of the form log(P(survival)/(1 − P(survival))) = β0 + β1(genotype) + β₂(cardenolide) + β₃(genotype × cardenolide). Post-hoc analysis tested whether individual genotype slopes (rate of change in survival with cardenolide concentration) differed significantly from zero to identify dose-dependent responses.

## Data availability

The data for this study are available on GitHub (https://github.com/mkarageorgi/MonarchFlies_Milkweeds).

## Code availability

The associated code for this study is available on GitHub (https://github.com/mkarageorgi/MonarchFlies_Milkweeds).

## Acknowledgements

This project was supported by grants from the National Institute of General Medical Sciences of the National Institutes of Health (Award number 1K99GM143455-01 to M.K, Award number R35GM119816 to N.K.W.), CalTeach Summer Research Institute grant to N.C.W, and a UC Berkeley Summer Undergraduate Research Program grant to E.O. The authors thank Christina Wistrom for her guidance to use greenhouse space at the Oxford Research Facility at UC Berkeley. The authors thank Alison Cutts for preparing milkweed graphics. The authors thank Niels Simon Groen, Rajeev Roy, and John Vontas for feedback that improved the manuscript.

## Author Contributions Statement

M.K., N.K.W. and A.A. designed research; M.K., N.C.W., M.A., D.Y., E.O., K.V., and S.L.B. performed research and were responsible for data acquisition; M.K. and N.C.W. analyzed data; M.K., N.C.W., A.A. and N.K.W. interpreted data; M.K. and N.C.W. wrote the first draft of the manuscript; M.K. prepared the figures; M.K., N.C.W., A.A. and N.K.W. revised the manuscript.

## Supplementary Figures

**Supplementary Figure 1.**
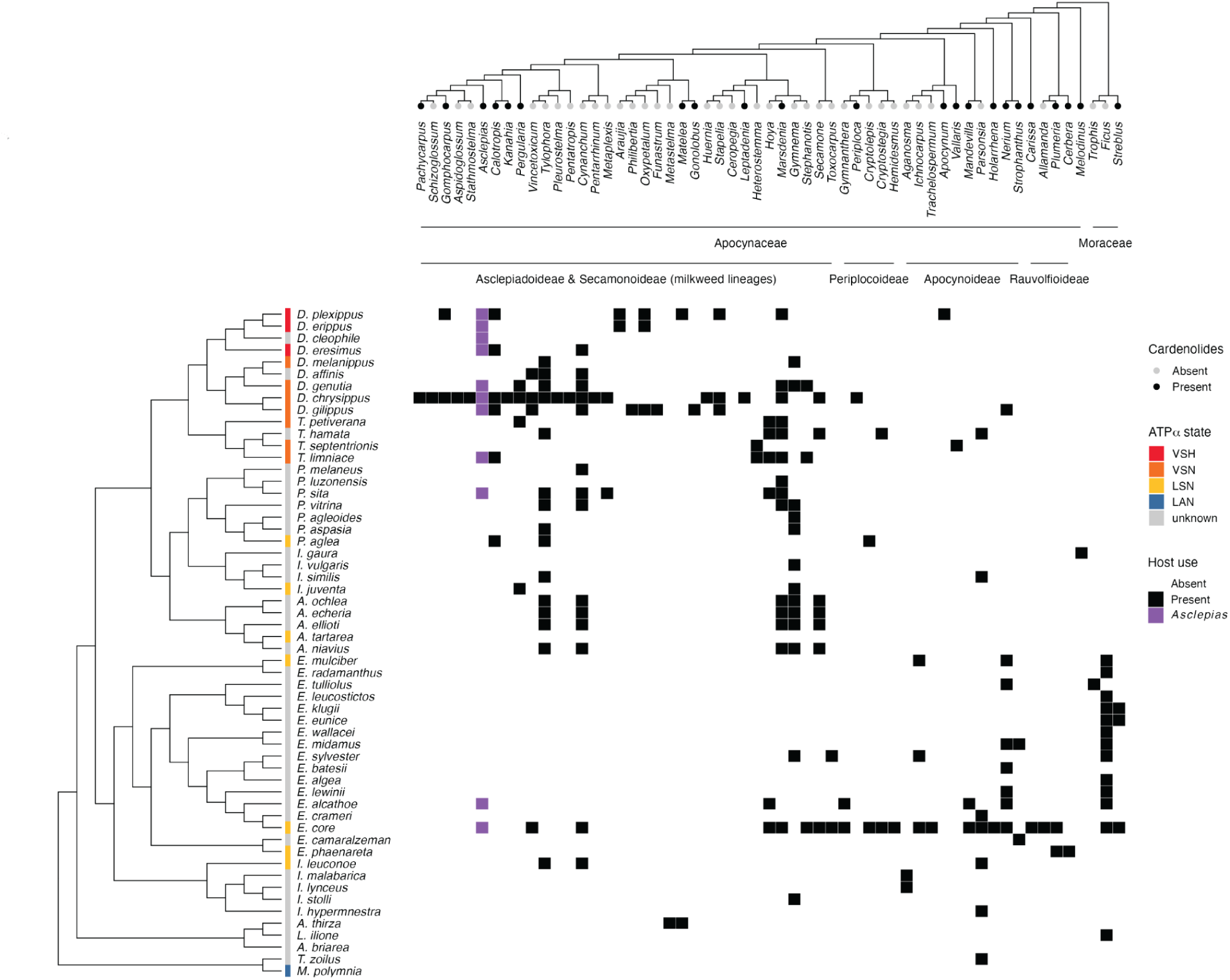
Host plant–butterfly matrix across 55 Danaini milkweed butterfly species and two outgroups within the Danainae subfamily. Rows and columns are ordered according to butterfly species and plant genus phylogenies, respectively. Cell shading indicates documented larval host-plant associations (black) or absence of recorded feeding (white) for each butterfly species–plant genus combination, with associations with the genus *Asclepias* highlighted in purple. Occurrence of cardenolides across plant genera is shown as top annotations. Phylogenies of Apocynaceae and Moraceae were grafted into a single composite tree. Six Apocynaceae genera present in historical host records were not sampled in the phylogeny of Fishbein et al. (2018) 29 and are therefore not represented in the matrix (see **Methods**).

**Supplementary Figure 2.**
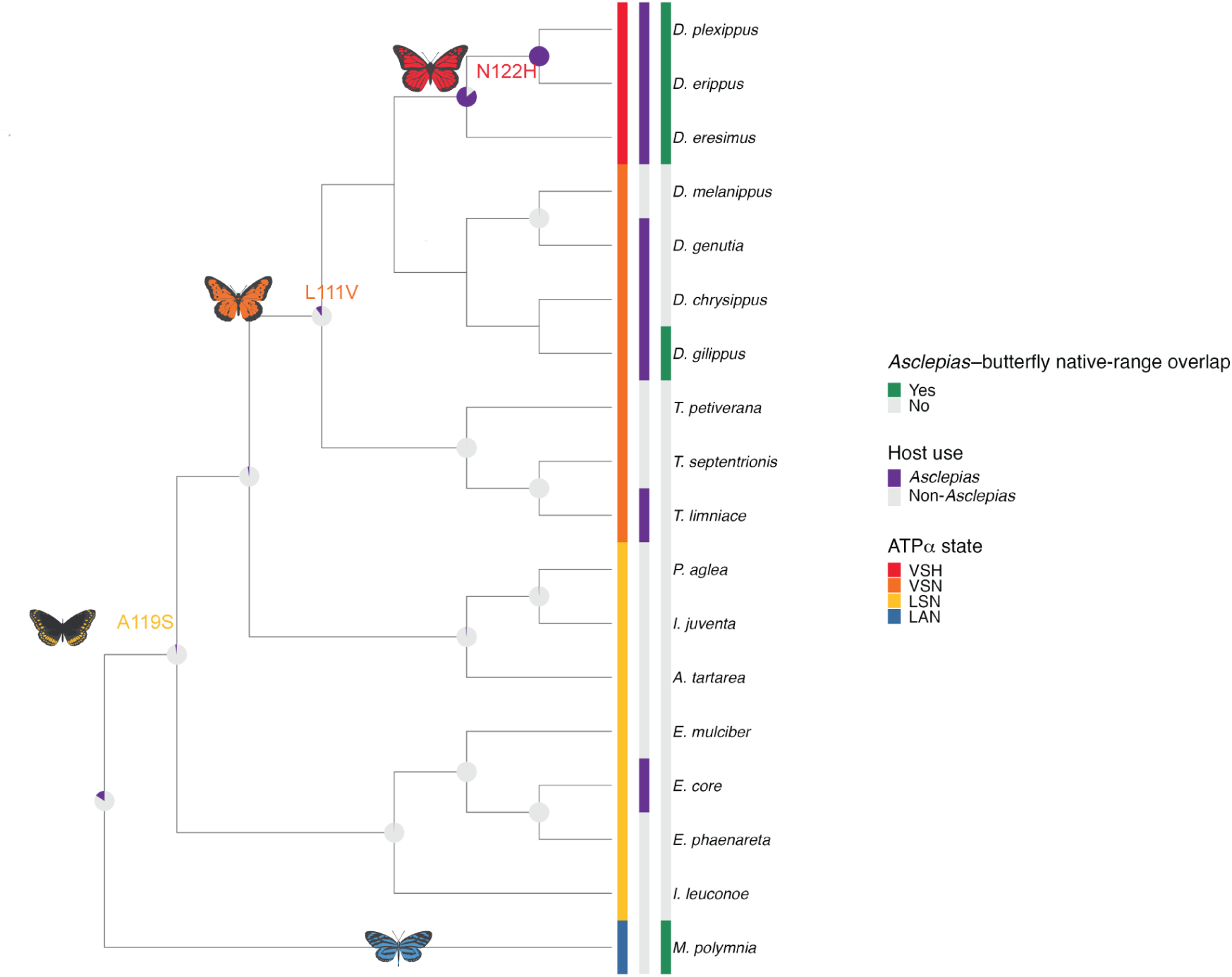
Geographically filtered reconstruction of native-range *Asclepias* association. Focused phylogeny of Danaini butterflies and one outgroup species with known ATPα states, corresponding to the subset shown in Fig. 1b. Pie charts show maximum-likelihood marginal probabilities of native-range *Asclepias* association, scored as present only when documented larval *Asclepias* use occurred in species whose native range overlaps the native range of *Asclepias* in the Americas (see **Methods**). Adjacent heatmaps show ATPα state, documented *Asclepias* host use, and native-range overlap with *Asclepias*. Documented *Asclepias* records outside this overlap were retained in the host-use dataset but scored as non-native-range associations in this geographically filtered reconstruction.

